# Phenethylaminylation: Preliminary *In Vitro* Evidence for the Covalent Transamidation of Psychedelic Phenethylamines to Glial Proteins using 3,5-Dimethoxy-4-(2-Propynyloxy)-Phenethylamine as a Model Compound

**DOI:** 10.1101/2025.02.13.638188

**Authors:** Rajiv S. Rangan, R. Max Petty, Suchismita Acharya, Kyle A. Emmitte, Rafael S. do Valle, Chandra Lam, Salman I. Essajee, William Mayhew, Olivia Young, Calvin D. Brooks, Michael J. Forster, Tara Tovar-Vidales, Abbot F. Clark

## Abstract

Psychedelics are well known for their ability to produce profoundly altered states of consciousness. But, more importantly, the effects of psychedelics can influence neurobehavioral changes that last well after these acute subjective effects end. This phenomenon is currently being leveraged in the development of psychedelic-assisted psychotherapies for the treatment of multiple neuropsychiatric disorders. The cellular and molecular mechanisms by which single doses of psychedelics are able to mediate long-term cognitive changes are an active area of research. We hypothesize that psychedelics contribute to long term changes in cellular state by covalently modifying proteins. This post-translational modification by psychedelics is possible through the transglutaminase-mediated transamidation of their amine termini to glutamine carboxamide residues. Here, we synthesize and utilize a propargylated analogue of mescaline – the classic serotonergic psychedelic phenethylamine found in cacti species – to identify putative protein targets of psychedelic modifications through the use of click-chemistry in a primary human astrocyte cell culture model. Our preliminary findings indicate that a diverse array of glial proteins may be substrates for transglutaminase 2-mediated monoaminylation by our model phenethylamine (“phenethylaminylation”). Based on these points, we speculatively highlight new directions for the study of this putative noncanonical psychedelic activity.

## Introduction

Psychedelics are a diverse group of phenomenologically unique, psychoactive chemical compounds known for their ability to produce profoundly altered states of consciousness^1–4^. Classic psychedelics include tryptamines (psilocybin/psilocin, *N,N*-dimethyltryptamine), lysergamides (lysergic acid diethylamide) and phenethylamines (mescaline), all of which exert their subjective effects through agonism of the serotonin (5-hydroxytryptamine, 5-HT) 2A receptor (5-HT_2A_)^5, 6^. Historically, cultural use of psychedelics has centered around their perceived ability to enable internal exploration of the mind, spiritual enlightenment, and healing^7–11^. In the last few decades, robust research has demonstrated the therapeutic potential of psychedelics in the treatment of an array of neuropsychiatric disorders, including anxiety, depression, post-traumatic stress disorder and substance use disorders ^12–17^. The value of psychedelic compounds as novel medicines lies in their low risk for adverse physiological effects, low abuse potential, and fast-acting – but enduring – therapeutic effects^18, 19^. For instance, “mystical-type” experiences brought upon by a single high-dose of psilocybin, a pro-drug of psilocin found in the polyphyletic fungal group known as “magic mushrooms,” can produce a sustained decrease in depressed mood and anxiety in cancer patients for at least six months^20–22^. The mechanisms by which psychedelics produce such long-lasting effects constitute an active area of study. A leading hypothesis in this area is psychedelic-induced neuroplasticity.

Neuroplasticity, the central nervous system’s (CNS) dynamic ability to adapt through the reorganization of synaptic connections, is involved in several higher-order neurological functions^23, 24^. The evidence that single doses of psychedelics can modulate neuroplasticity and thereby produce persistent neurobehavioral effects has been reviewed thoroughly elsewhere^25,26^. In brief, psychedelic-activated signaling downstream of 5-HT_2A_ leads to structural and functional neuroplasticity. This results in an acute period of increased synaptic growth and strengthened long-term potentiation, especially in cortical regions of the brain with a high density of 5-HT_2A_. The neural rewiring that occurs during this window of heightened neuroplasticity is likely durable and may be responsible for the permanence of effects associated with single doses of psychedelics^25, 26^. However, the molecular mechanisms that underpin these responses to psychedelics by cells of the CNS are still not fully understood.

Interestingly, an *in vitro* study previously demonstrated that 5-HT_2A_ activation by 2,5-dimethoxy-4-iodoamphetamine (DOI; a substituted amphetamine and psychedelic phenethylamine) transiently increases dendritic spine size in primary cortical neurons. This effect was, in part, dependent on the transamidation of the small GTPases Rac1 and Cdc42 by the enzyme transglutaminase 2 (TGM2)^27^. TGM2 canonically carries out the calcium-dependent transamidation of lysine ε-amino groups to glutamine γ-carboxamide groups, enabling it to covalently crosslink proteins^28–31^. TGM2 possesses numerous diverse functions, however, including the ability to covalently attach biogenic monoamines (e.g., dopamine and serotonin) to proteins by catalyzing the transamidation of their primary amine termini to the amide of available glutamine residues on various protein targets. This post-translational modification is broadly referred to as monoaminylation, or specifically as dopaminylation, serotonylation, and so forth^32–36^. Monoaminylation modulates the localization, state, and function of a variety of proteins^36–40^. In this manner, monoaminylation can act as an atypical signaling pathway by which monoamine neurotransmitters exert their effects throughout the CNS^41, 42^. Of particular interest, serotonylation and dopaminylation modify the glutamine 5 residue on histone H3 proteins (H3Q5). Serotonylation of histone H3 tri-methylated lysine 4 (H3K4me3)-marked nucleosomes, in particular, appears to be a transcriptionally permissive modification, altering the interaction of reader proteins with H3 and modulating the expression of numerous genes^43–50^. In neurons, this type of histone monoaminylation is a potentially stable modification, possibly allowing monoamine neurotransmitters to directly modulate long-term changes in chromatin organization and gene expression^51–54^. Given the interaction between psychedelic signaling pathways and TGM2, it is plausible that the mediation of histone (and other protein) modifications by monoamines may be a mechanism by which psychedelics promote chronic neurobiological changes.

We hypothesize that a subset of chemically amenable psychedelics may directly modify protein targets. Exogenous psychedelic tryptamines and phenethylamines are structurally similar to endogenous monoamine neurotransmitters (Fig. 1). Some psychedelics, such as mescaline (a phenethylamine produced in cacti such as peyote, *Lophophora williamsii*) have primary amine groups on their amine termini and should be readily transamidated onto glutamine γ-carboxamide groups^55–57^. Others, such as norpsilocin and psilocin have secondary and tertiary amine termini, respectively^58–61^. These compounds *in vivo* encounter demethylases and, thereby, may also act as substrates for TGM2^62, 63^. Although it is unclear if demethylation could occur at a rate sufficient to produce meaningful quantities of monoaminylation substrates, given the presence of competing psychedelic metabolism pathways^64^. Such a mechanism could be useful in explaining specific therapeutic effects of psychedelics, including their possible antidepressant and anti-addictive qualities^65, 66^. Serotonylation of H3K4me3 in the dorsal raphe is dysregulated by stress. Remarkably, serotonergic antidepressants can disrupt H3K4me3 serotonylation, leading to an attenuation of stress-related gene expression^67^. Dopaminylation of H3 in the ventral tegmental area is increased in rats experiencing cocaine withdrawal. Disruption of H3 dopaminylation attenuates cocaine-related gene expression and reduces cocaine-seeking behavior^68^. Perhaps competing modifications at H3Q5 by psychedelics could promote a return to homeostatic chromatin states in these critical brain regions.

**Figure 1.**
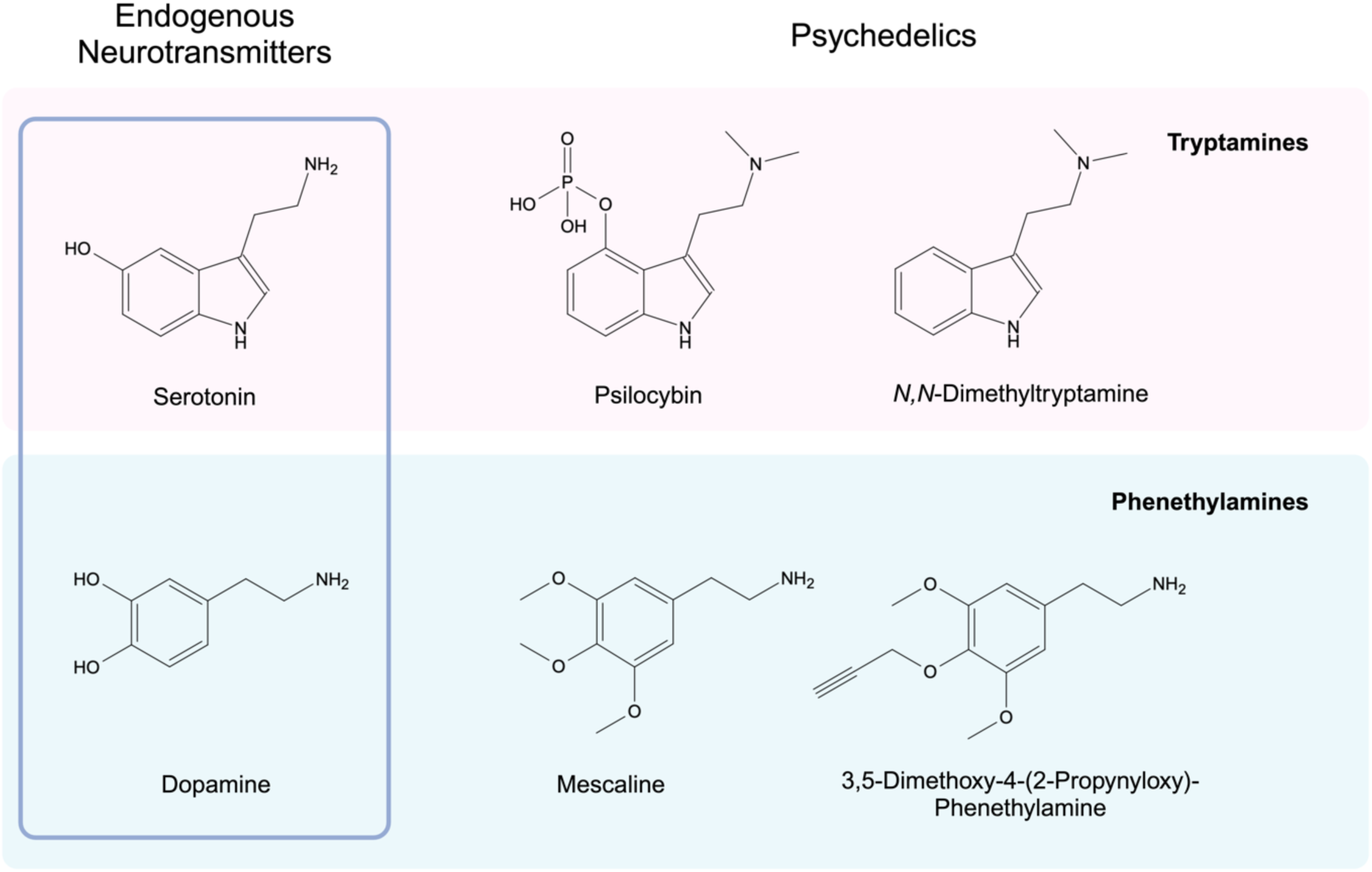
Psychedelic tryptamines and phenethylamines are structurally similar to the endogenous neurotransmitters serotonin and dopamine, respectively.

The idea that psychoactive tryptamines, phenethylamines, and amphetamines may act as substrates for TGM2-mediated monoaminylation has been considered by other groups. In a recent review, Amni Al-Kachak & Ian Maze proposed the possibility of “amphetaminylation”^69^. In another, Joseph Benetatos provides a thoughtful examination of the hypothesis of psychedelic protein modifications and its potential implications^70^. However, covalent protein modification by a psychedelic compound has not yet been experimentally demonstrated. This may be due to the fact that exotic protein modifications are difficult to detect^42^. In the case of serotonylation, one solution to enable the enrichment and detection of serotonylated peptides is the use of a alkyne-functionalized derivative of serotonin, 5-propargloxytryptamine (5PT)^71^. The presence of a terminal alkyne on 5PT enables the use of “click-chemistry” enrichment techniques. For example, the Copper(I)-catalyzed Azide-Alkyne Cycloaddition (CuAAC) can be used to attach a biotinylated azide to proteins “serotonylated” with 5PT. Biotin pulldown assays can then be used to enrich all modified proteins^48, 72^. In this study, we utilize a similar technique to provide an initial identification of potential protein targets of TGM2-mediated modification with exogenous psychedelic monoamines.

In cell culture models, mescaline serves as an ideal candidate for preliminary investigations into the possibility of protein modification by psychedelics given its primary amine terminus. Conveniently, an alkyne-functionalized derivative of mescaline has previously been described. 3,5-dimethoxy-4-(2-propynyloxy)-phenethylamine was first synthesized and described by Alexander Shulgin in his landmark work *PiHKAL* (#143, “Propynyl”)^73^. 3,5-dimethoxy-4-(2-propynyloxy)-phenethylamine shares the same basic chemical structure as mescaline, with a terminal alkyne present on the 4-methoxy group (Fig. 1). This enables it to be used in place of mescaline for identifying proteins monoaminylated by such psychedelic phenethylamines (“phenethylaminylation”), with the advantage of being amenable to azide-alkyne click-chemistry and subsequent enrichment via biotin pulldown (Fig. 2). In our study, we propose a simplified synthetic route for the production of 3,5-dimethoxy-4-(2-propnyloxy)-phenethylamine and demonstrate its potential usefulness for detecting proteins modified by psychedelic phenethylamines via click-chemistry in a primary human astrocyte cell culture model.

**Figure 2.**
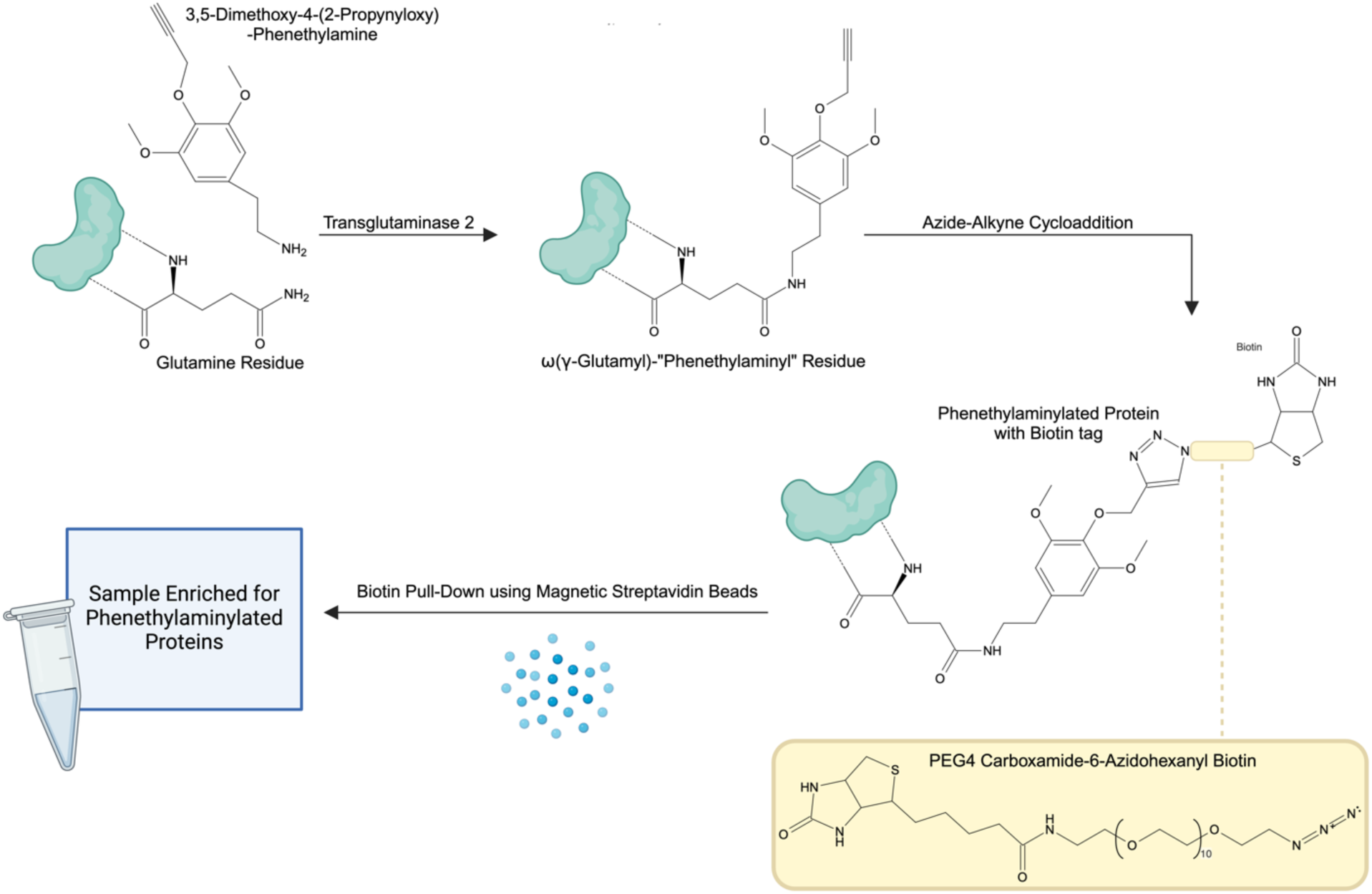
Method for the enrichment of proteins phenethylaminylated by 3,5-dimethoxy-4-(2-propynyloxy)-phenethylamine. Residue structure depiction and nomenclature inspired by work of Michael Bader^42^.

## Chemistry

### Synthesis of 3,5-Dimethoxy-4-(2-Propynyloxy)-Phenethylamine

In the first step, 3,5-dimethoxy-4-(2-propynyloxy)-benzaldehyde was synthesized from 4-hydroxy-3,5-dimethoxybenzaldehyde (syringaldehyde, commercially available; Cayman Chemical, Ann Arbor, MI) using a method described by Razzano, et al. (Fig. 3a)^74^. A mixture of 1.0 g (5.5 mmol, 1.0 eq) syringaldehyde, 2.3 g (16.6 mmol, 3.0 eq) K_2_CO_3_, and 1.9 mL (80% in toluene, 21.3 mmol, 3.9 eq) propargyl bromide in acetonitrile was refluxed for 3 hours under a N_2_ atmosphere. The precipitate was then filtered out, and the filtrate was concentrated via rotary evaporation under reduced pressure (rotavap). The solids were dissolved in ethyl acetate and partitioned with water. The organic phase was washed with a saturated NaCl solution (brine) and dried over sodium sulphate before being concentrated by rotavap. The solids were dissolved in dichloromethane and flash purification was performed on a Biotage Isolera Prime system using a mobile phase of hexane with increasing %ethyl acetate, producing 0.89 g (4.0 mmol, 73%) of 3,5-dimethoxy-4-(2-propynyloxy)-benzaldehyde as a white powder that was carried forward into the next step.

**Figure 3.**
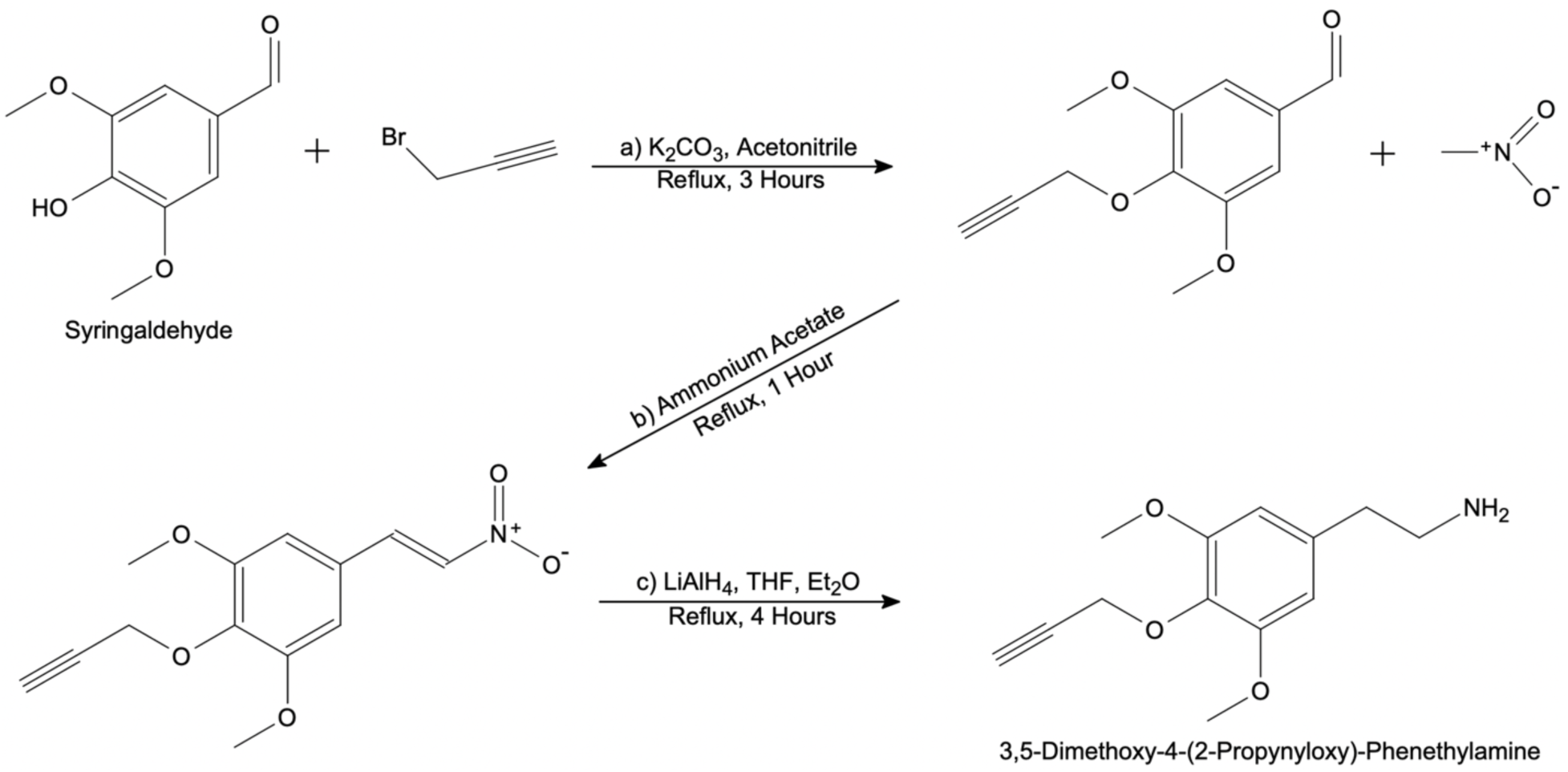
Synthetic scheme for the preparation of 3,5-dimethoxy-4-(2-propynyloxy)-phenethylamine in three steps: a) propargylation, b) nitration, c) reduction.

In the second step, 3,5-dimethoxy-4-(2-propynyloxy)-nitrostyrene was synthesized from 3,5-dimethoxy-4-(2-propynyloxy)-benzaldehyde using a method described by Ding, et al. (Fig. 3b)^75^. A mixture of 1.0 g (4.5 mmol, 1.0 eq) 3,5-dimethoxy-4-(2-propynyloxy)-benzaldehyde, 9.0 mL (169.0 mmol, 37.3 eq) nitromethane, and 0.52 g (6.8 mmol, 1.5 eq) ammonium acetate was refluxed for 1 hour under a N_2_ atmosphere. The mixture was then concentrated by rotavap, producing a reddish brown/orange residue. The residue was recrystallized by first fully dissolving in hot ethanol and then rapidly cooling on ice, yielding 0.46 g (2.1 mmol, 48%) of fine yellow needles.

In the third step, 3,5-dimethoxy-4-(2-propynyloxy)-nitrostyrene was reduced to 3,5-dimethoxy-4-(2-propynyloxy)-phenethylamine using a method modified from the procedures described by Ding, et al. and Shulgin (Fig. 3c)^73, 75^. 0.43 g (2.0 mmol, 1.0 eq) of the product from step 2 was dissolved in anhydrous tetrahydrofuran (THF) and added dropwise to a well-stirring suspension of 0.31 g (8.2 mmol, 5.0 eq) LiAlH_4_ in diethyl ether (Et_2_O) under N_2_ at 0 °C. The neat reaction mixture was refluxed for 4 hours under a N_2_ atmosphere and then cooled to 0 °C. Isopropyl alcohol was slowly added to the mixture to quench the reaction. An equal volume of 3.75 M NaOH was then slowly added with gentle swirling. The mixture was brought up to room temperature and stirred for 30 minutes, producing slightly tan solids. Solids were filtered out and the filter cake was washed with THF. The filtrate was concentrated by rotavap, producing an oily yellow residue. The residue was dissolved in 0.5 M H_2_SO_4_. In a separatory funnel, this aqueous solution was washed twice with dichloromethane. The aqueous phase was then made basic using 1 M NaOH and partitioned using dichloromethane. The organic phase was extracted and dried over sodium sulphate, before being concentrated by rotavap, again producing an oily yellow residue. The residue was dissolved in isopropyl alcohol, acidified with concentrated HCl and then diluted with Et_2_O. Solids precipitated out of solution and were filtered and washed with Et_2_O, yielding 40 mg (0.15 mmol, 7.5%) of 3,5-dimethoxy-4-(2-propynyloxy)-phenethylamine hydrochloride as an off-white powder. Upon dissolution in water followed by lyophilization, characteristic white needles appeared^73^.

All reagents used in the above chemical synthesis were ACS reagent grade. Yield was not optimized for this study. The identity of 3,5-dimethoxy-4-(2-propynyloxy)-phenethylamine was confirmed using ^1^H NMR, ^13^C NMR, and ToF mass spectrometry (Fig. 4). ^1^H NMR (300 MHz, D_2_O): δ 2.76 ppm (t, *J* = 2.46 Hz, 1H), 2.86 ppm (t, *J* = 7.16 Hz, 2H), 3.17 ppm (t, *J* = 7.24 Hz, 2H), 3.76 ppm (s, 6H), 4.62 ppm (d, *J* = 2.39, 2H), 6.61 ppm (s, 2H). ^13^C NMR (300 MHz, D_2_O): δ 33.01, 40.43, 56.02, 59.96, 76.41, 78.89, 106.16, 132.95, 134.21, 152.97 ppm. MS (LC-ToF, H_2_O): predicted *m/z* for ([C_13_H_17_NO_3_]+H)+ = 236.1281, determined *m/z* = 236.1279. Purity was estimated to be >97% based on HPLC at 260 nm.

**Figure 4:**
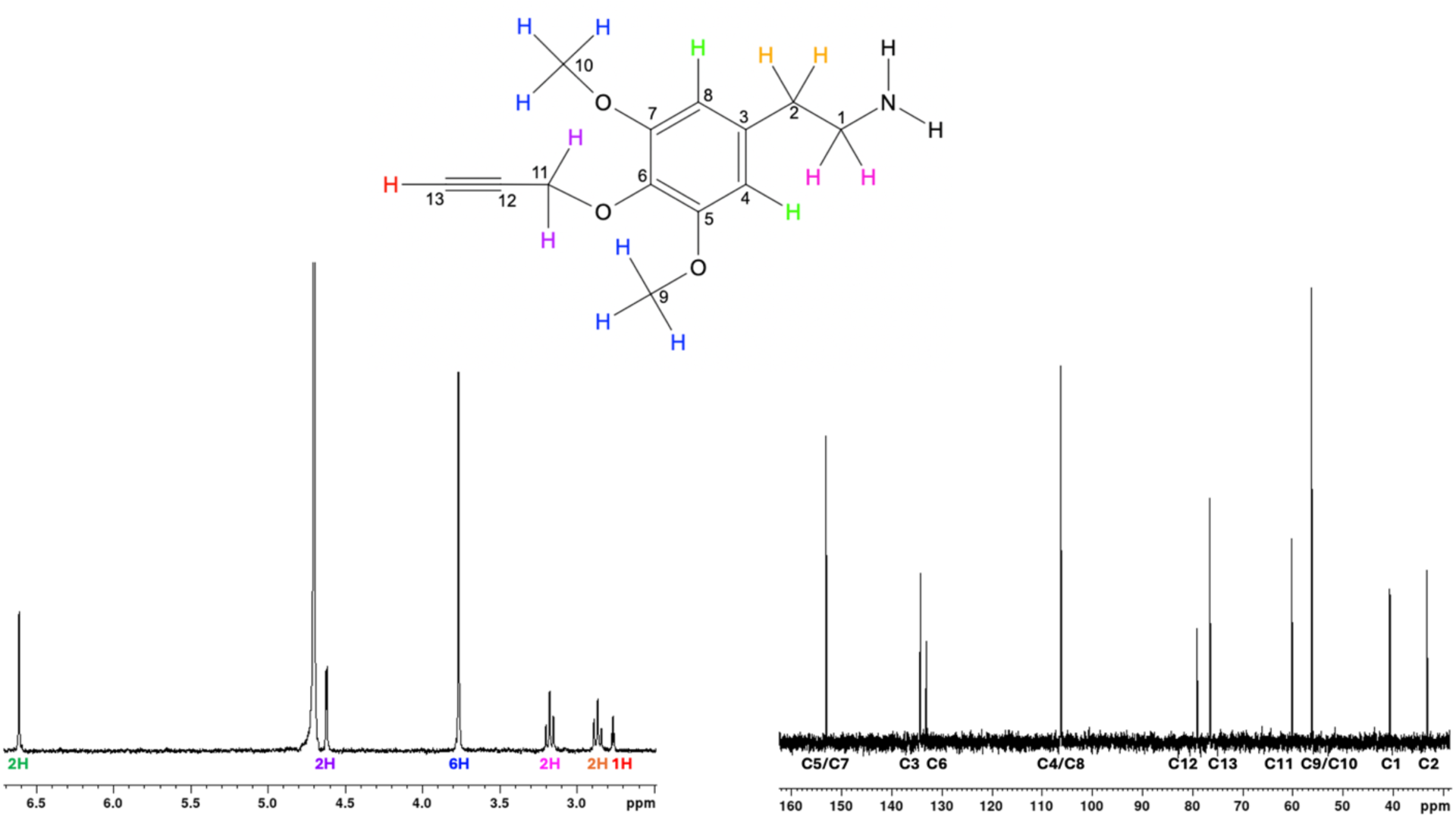
3,5-Dimethoxy-4-(2-Propynyloxy)-Phenethylamine structure validation by ^1^H NMR (left, with protons color-coded) and ^13^C NMR (right, with carbons numbered).

For simplicity, we will refer to 3,5-dimethoxy-4-(2-propynyloxy)-phenethylamine as “4-propargylmescaline” or “4PM” during the following discussion of our results.

## Results

### Cell Permeability of 4-Propargylmescaline

Although TGM2 localizes to multiple cellular compartments and the extracellular milieu, it is generally thought to act as a transamidase primarily in the intracellular space^76, 77^. As such, for 4PM to function as a probe for TGM2-mediated phenethylaminylation, it must be taken up by cells. To assess the uptake of 4PM by ONH astrocytes and determine doses at which its intracellular presence can be easily detected, we performed click-chemistry on fixed cells to biotin-tag 4PM molecules for fluorescence detection by AlexaFluor-647 fluorophore-conjugated streptavidin.

After click-chemistry, we were able to observe intracellular 4PM via fluorescence microscopy (Fig. 5A). Compared to untreated controls, fluorescence intensity was significantly increased in the 100 μM (15.82-fold, p=0.015), 500 μM (38.22-fold, p=0.041) and 1000 μM (41.00-fold, p=0.046) treatment groups (Fig. 5B). Imaging parameters were set identically across groups and were optimized to avoid oversaturation of fluorescence signal in the 1000 μM treatment group. As such, signal from the 50 μM was more difficult to detect. Our rational for this approach was to ensure that our selected dose was sufficient to elicit obvious effects in treated cells, given the preliminary nature of our experiments. Based on these findings, a 100 μM concentration of 4PM was selected for use in the following experiments.

**Figure 5.**
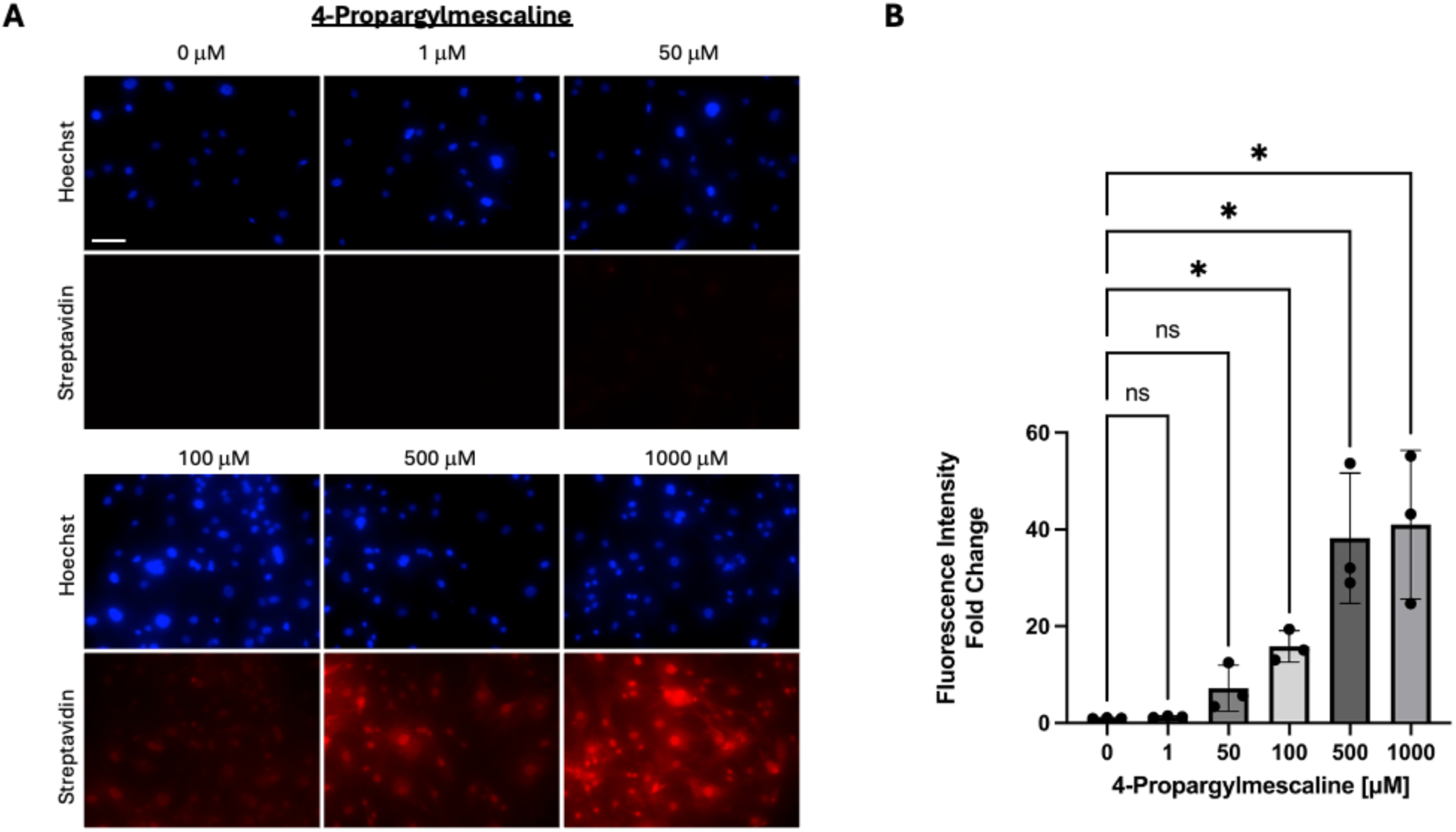
Detection and quantification of intracellular 4PM in ONH astrocytes. A) Streptavidin-mediated detection of 4PM molecules “clicked” to biotin is represented by fluorescent signal (red). Total magnification = 200x, scale bar = 100 μm (first panel). B) Fluorescence intensity analysis with values represented as fold-changes relative to average control fluorescence intensity (*indicates p<0.05, biological n=3, error bars represent SD).

**Figure 6.**
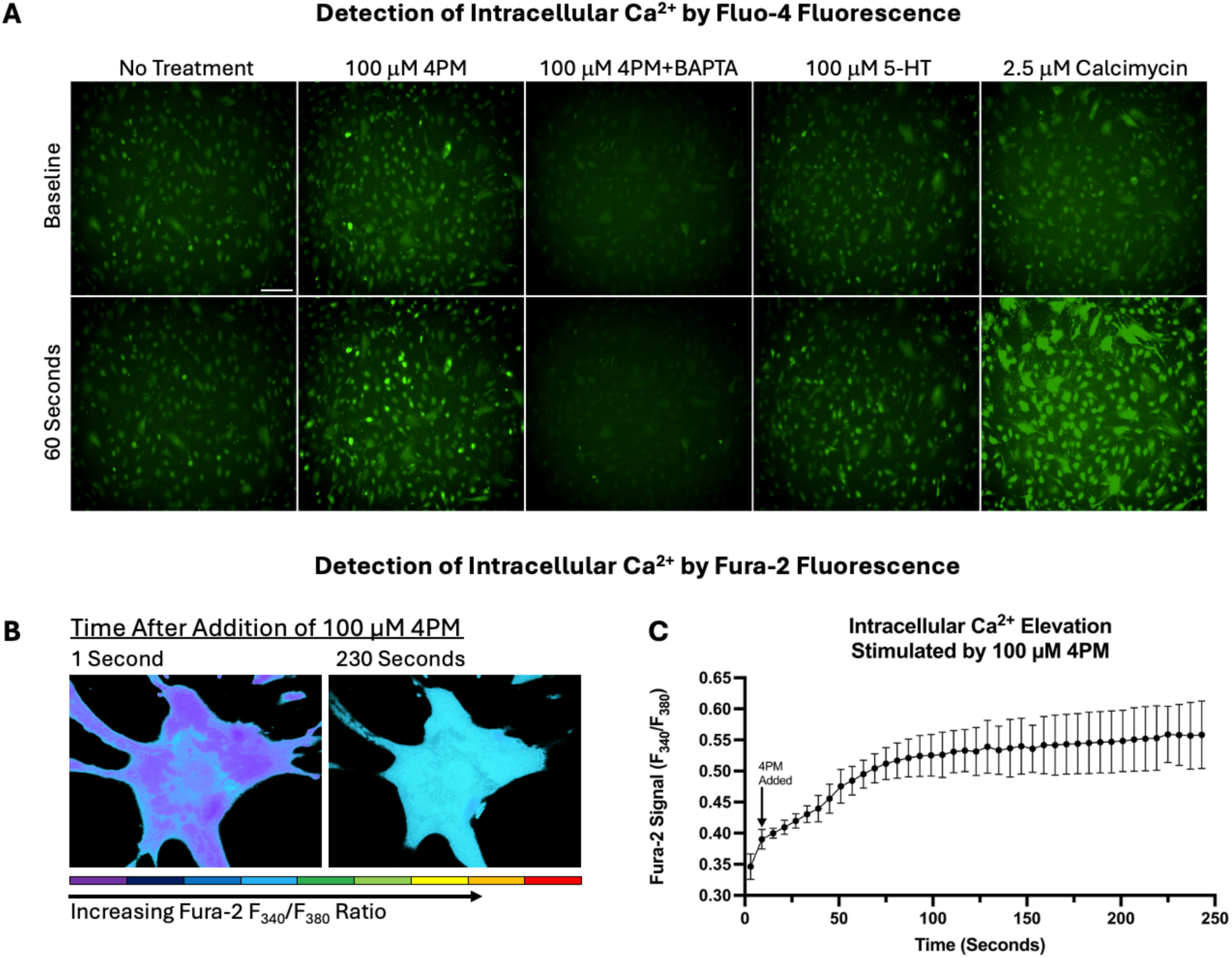
Fluorescent analysis of intracellular Ca^2+^ elevation induced by 100 μM 4PM. A) Representative images demonstrating increased green fluorescence of the calcium indicator Fluo-4 in ONH astrocytes 1 minute after the addition of 100 μM 4PM, 100 μM 5-HT or 2.5 μM Calcimycin (ionophore, positive control). Fluo-4 signal is ablated in the presence of 10 μM BAPTA, a calcium chelator. Total magnification = 100x, scale bar = 100 μm (first panel). B) Representative pseudocolored images demonstrating changes in intracellular Ca^2+^ level/Fura-2 signal in a single ONH astrocyte between 1 second after the addition of 100 μM 4PM and 230 seconds after the addition of 100 μM 4PM (peak observed effect). Cooler colors represent lower Fura-2/calcium signal (F_340_/F_380_); warmer colors represent higher signal. Total magnification = 400x. C) Quantified Fura-2 ratio (F_340_/F_380_) plotted against time after the addition of 100 μM 4PM. Error bars represent standard deviation among biological replicates (n=3).

The apparent permeability of cultured cells to a mescaline-like compound at high doses is suspected to be partly due to internalization via various organic cation transporters, although this is not tested^78–80^.

### 4-Propargylmescaline Stimulates Modest Elevation of Intracellular Calcium

The transamidation activity of TGM2 is calcium ion (Ca^2+^)-dependent^81^. Previous studies have demonstrated that serotonin and serotonergic 5-HT_2A_ agonists are able to induce TGM2-mediated protein cross-linking and protein monoaminylation through downstream calcium signaling^32, 36, 82–84^. To determine if 4PM could similarly stimulate calcium signaling and thereby reasonably be expected to promote TGM2 activity, we assessed the ability of 4PM to elevate intracellular Ca^2+^ levels in ONH astrocytes via a qualitative Fluo-4 assay and a quantitative Fura-2 assay. 100 μm 4PM produced a modest increase in fluorescent signal from Fluo-4, which was attenuated by the addition of a calcium chelator (BAPTA), indicating an increase in intracellular Ca^2+^ available for binding by the indicator. This Ca^2+^ increase was qualitatively similar to the effect observed with 100 μm serotonin (5-HT). Using the Fura-2 F_340_/F_380_ fluorescence ratio, we again observed that 100 μm 4PM produced a modest (∼1.60-fold) intracellular Ca^2+^ elevation, peaking between 2-3 minutes.

Many experiments have demonstrated evidence for transamidation modifications of cellular proteins (including monoaminylation) in cultured cells under conditions in which large Ca^2+^ elevations are not likely expected ^33, 34, 85–87^. These observations may be due to the fact that in living cells TGM2 can have variable sensitivities to activating or inhibiting stimuli^88^. For instance, a shorter TGM2 isoform expressed in astrocytes is less sensitive to inhibition by GTP and is, resultingly, more prone to Ca^2+^ activation^89^. Despite these potential exceptions, TGM2 is generally thought to be latent in the absence of substantial intracellular Ca^2+^ elevations^90^. TGM2 also likely requires prolonged calcium elevation for robust activation^91^. Although our examination of the effects of 4PM on cellular Ca^2+^ dynamics is limited, our data indicate that 4PM alone may not stimulate a consistently robust elevation in TGM2 activity across all of our cell strains. For this reason, we opted to co-treat our cells with Calcimycin in subsequent experiments to ensure that sufficient Ca^2+^ is available for potential phenethylaminylation to occur – this approach has been previously used to stimulate the TGM2-mediated incorporation of histamine onto proteins *in vitro*^92^.

### Identification of Potential Phenethylaminylation Protein Targets

Samples enriched for phenethylaminylated proteins were prepared by streptavidin-magnetic bead pulldown after biotin tags were “clicked” onto proteins modified by 4PM. On an immunoblot probed for biotin (Fig. 7), we observed a clear enrichment of biotinylated protein bands (∼1.74-fold, p=0.048). This effect is prominent at several medium molecular weight bands and on one <15 kDa band. A strong nonspecific signal is seen at ∼15 kDa due to leeching of streptavidin monomers. Despite efforts to reduce the presence of nonspecific sources of biotinylation, much of the “basal” biotin signal seen in the vehicle group is suspected to be the result of residual endogenous biotinylation, nonspecific binding of the biotin-azide probe and nonspecific antibody signal. In samples from cells pre-treated with the irreversible, selective TGM2 inhibitor Z-DON, this enriched biotin signal is reduced by ∼1.29-fold (p=0.020) (Fig. 7), suggesting that the biotin signal associated with 4PM is likely to be a consequence of TGM2-mediated transamidation/phenethylaminylation. This analysis should be interpreted as only semi-quantitative, however, as band intensity is variable and some bands fall outside the linear range of detection. The incomplete inhibition (∼53%) of TGM2-mediated 4PM-biotin signal by Z-DON is partly explainable by the fact that some non-enzymatic cysteine labeling by monoamines is expected to occur *in cellulo*^93, 94^.

**Figure 7.**
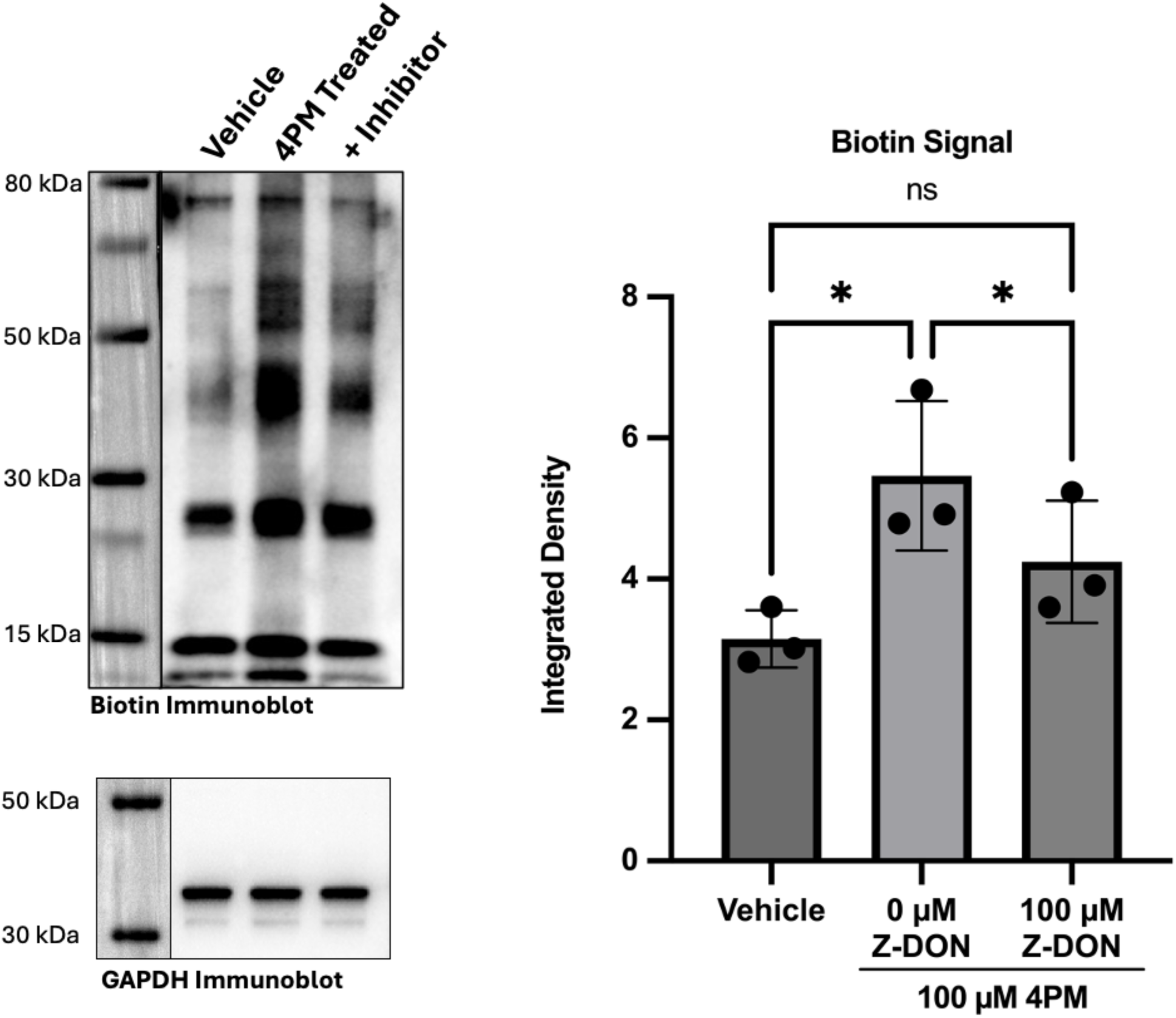
Representative biotin immunoblot demonstrating enrichment of biotinylated proteins – phenethylaminylated proteins modified by 4PM “clicked” with biotin tags – after streptavidin:biotin pulldown. Experimental groups were treated with a vehicle solution, 100 μM 4PM, or 100 μM 4PM + 100 μM Z-DON, respectively. A semi-quantitative integrated density analysis comparing the total biotin signal normalized to input GAPDH for the groups is provided (*indicates p<0.05, biological n=3, error bars represent SD).

Given the observation of biotinylated protein enrichment after click-chemistry/biotinylation and streptavidin pulldown, we moved forward with LC-MS identification of potentially phenethylaminylated proteins in 4PM-treated samples. By examining the most abundant proteins in our enriched samples (Fig. 8), we hope to highlight potential targets of phenethylaminylation, although some of these abundant proteins certainly represent nonspecific capture^95^. While our data cannot be used to confirm any protein modifications, it does suggest that a variety of substrates could be targets of phenethylaminylation, as is true for monoaminylation broadly. These targets could include extracellular matrix proteins, cytoskeletal proteins, metabolic enzymes, small G-proteins, heat shock proteins, ribosomal proteins, among others.

**Figure 8.**
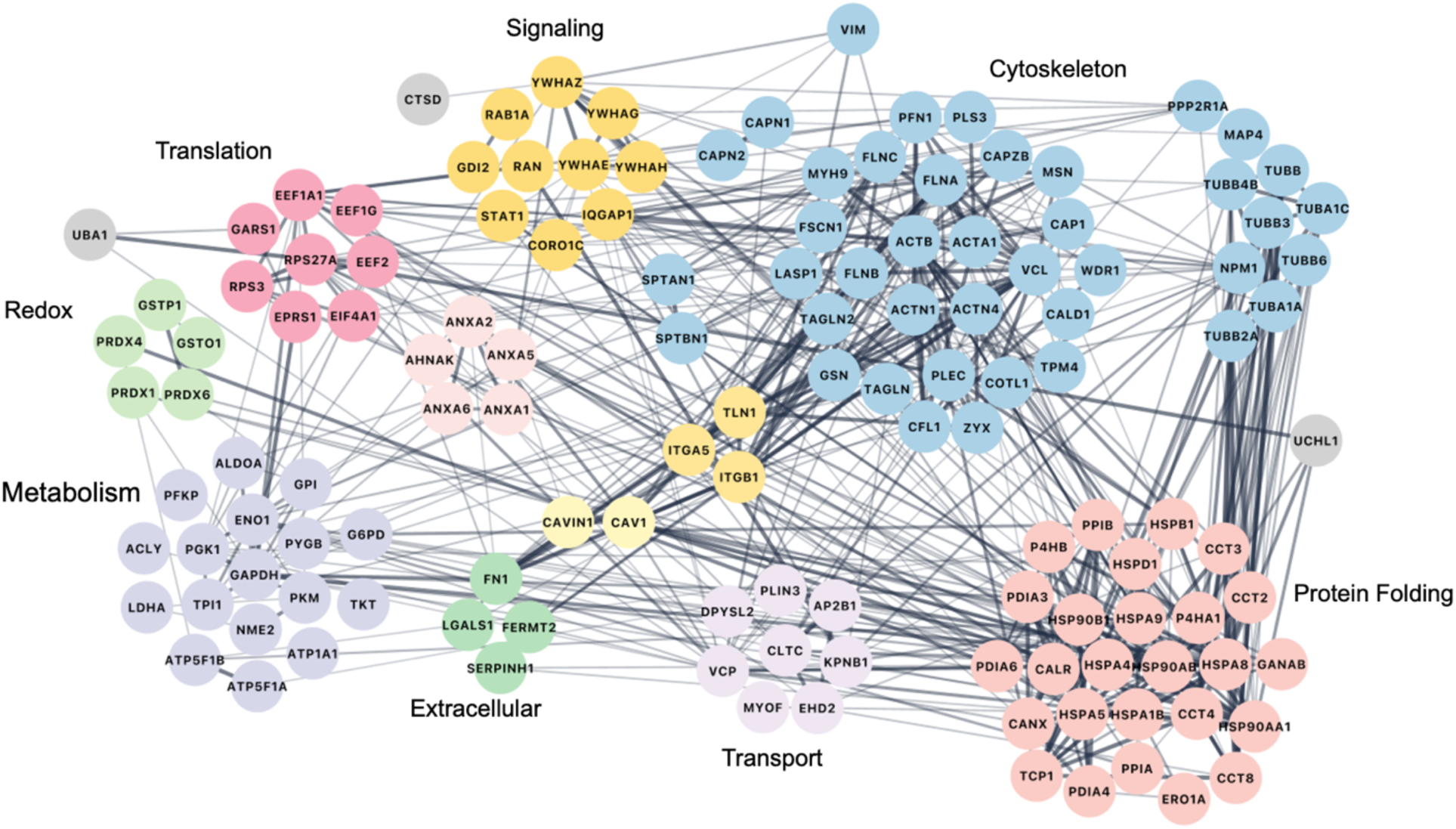
String physical interaction network for potentially phenethylaminylated proteins – the most abundant proteins identified in 4PM-treated samples after biotinylation via click-chemistry and streptavidin:biotin pulldown.

## Discussion

Our findings provide a preliminary indication that the TGM2-catalzyed transamidation of a phenethylamine structurally similar to the classic psychedelic mescaline may be able to covalently modify a broad range of protein substrates. We believe that this activity is likely to occur with other psychoactive compounds whose pharmacophores contain amine termini – phenethylamines, amphetamines, tryptamines.

These results must be considered in the light of several substantial limitations. First, our study was conducted using supraphysiologic conditions including high compound doses, long treatment times, and artificially elevated intracellular Ca^2+^ levels. Physiologically meaningful doses are much lower than the 100 μM dose used in this study, based on previous reports of mescaline and DOI biodistribution in the brain^96, 97^. Moreover, mescaline, like other psychedelics, is rapidly cleared from the brain^97^. Further investigation will need to be conducted using physiologically relevant parameters, based on the biodistribution of 4PM *in vivo* at therapeutically relevant doses, to determine if phenethylaminylation is a biologically meaningful process. The physiologic relevance of modifications by 4PM is also complicated by the fact that it may not elicit subjective psychedelic effects comparable to those of mescaline or DOI, based on the limited reports of its consumption by humans^73^. Behavioral tests, such as the head-twitch response in murine models, may be useful in partially characterizing its effects during future studies^98, 99^. Such studies would also benefit from optimized synthetic methods for improved chemical yield and purity. Second, our findings in adult primary ONH astrocytes may not be translatable to other cell types of the CNS, such as neurons or protoplasmic, gray-matter astrocytes^100^. While we can speculate on the potential effects of specific phenethylaminylated proteins in various neural cells and tissues, future studies must be performed on directly relevant cell populations when investigating a particular physiologic, pathophysiologic, or therapeutic process. Yet, the use of primary human astrocytes does have intrinsic value. Glial cells are likely to be involved in the brain’s response to psychedelics and exert strong cellular responses upon 5-HT_2A_ activation^24, 101^. Given the role of glia, especially astrocytes, in the regulation of CNS metabolism, synapse development, cell-cell signaling and other homeostatic processes, the impact of canonical and noncanonical psychedelic signaling pathways in astrocytes is certainly understudied^102, 103^. Lastly, and most importantly, we do not characterize specific chemical adducts between our model phenethylamine and individual peptide sequences. For this reason, our data should not be viewed as definitive evidence that any particular protein is phenethylaminylated but should serve as a proof-of-concept that a broad range of protein substrates for phenethylaminylation likely exist. In future studies, robust biochemical validation will need to be employed to confirm the hypothetical modification described in this paper. After such experiments are conducted, studies can be performed to determine the functional consequences, if any, of monoaminylation by psychedelics at specific protein sites.

With these limitations in mind, our study highlights a potential new direction in the research of atypical psychedelic signaling pathways. We found that 4PM labeling was associated with cytoskeletal proteins, extracellular matrix proteins, small signaling molecules, heat shock proteins, metabolic enzymes, ribosome-associated proteins and more. These results are congruent with reports of monoaminylated protein targets in the broader literature, many of which have documented specific monoamine-modified glutamine sites^72, 104, 105^. Recently, Zhang et al. have robustly confirmed the dopaminylation and serotonylation of numerous proteins in cancer cells using new chemical labeling techniques^93, 106^. The diversity of peptides plausibly available for covalent modification by psychedelic monoamines represents the variety of novel mechanisms by which these drugs may modulate long term changes in cellular structure and function. In this discussion, we will briefly review the broader literature on monoaminylation in order to speculate on a few such plausible mechanisms.

### Actins

Serotonylation of actins and actin-associated proteins modulate cytoskeletal contractility in smooth muscle cells^40^. Interestingly, in a cortical cell line, DOI-stimulated 5-HT_2A_ activation increases transamidation of Rac1, which influences neuronal differentiation and neurite outgrowth through regulation of the actin cytoskeleton^82^. The transamidation of psychedelics to cytoskeletal proteins and their regulators may serve as a mechanism by which they influence the structure of synaptic connections among neurons^70^. The interaction of astrocytic processes with synapses is important in their regulation and plasticity^107^. The hypothetical modulation of actin dynamics by actin-psychedelic adducts in astrocytes themselves could also be important to the neuroplastic effects of psychedelics.

### GTPases

Serotonylation modifies numerous GTPases, including RhoA, Rab4, Rab3a, Rab27a, while histaminylation (monoaminylation by histamine) modifies Cdc42, G_αo1_ and G_αq_. Through these modifications, monoaminylation can influence an array of cellular processes related to G-protein coupled receptor (GPCR) signaling including intracellular calcium regulation, metabolism, endocytosis, exocytosis, and more^36, 84, 108–113^. Intriguingly, psychoactive compounds with activity at the serotonin transporter (SLC6A4) – fluoxetine and tramadol – may modulate levels of GTPase serotonylation^114, 115^. Monoaminylation of GTPases likely instigates constitutive activation^116^. Therefore, psychedelics, through the regulation of monoaminylation or through direct monoaminylation of GTPases, may mediate lasting modifications to cellular responses related to GPCR-neurotransmitter signaling in a manner similar to that of endogenous and other exogenous aminergic compounds^112^.

### Metabolic Enzymes

Serotonylation of glyceraldehyde-3-phosphate dehydrogenase (GAPDH) promotes its cytosolic localization. Dopaminylation of triosephosphate isomerase 1 (TPI1) promotes its enzymatic activity. In both cases, monoaminylation shifts cells towards glucose metabolism^37, 117^. Protein modification by psychedelics could add another layer to the observations of glucose metabolism modulation by *N,N*-dimethyltryptamine (DMT) and other psychedelics.^118, 119^.

### Heat Shock Proteins

Although the effects of monoaminylation on heat shock proteins have not yet been investigated, they are well documented as substrates for various TGM2 activities, including monoaminylation^72^. The interaction between heat shock proteins and TGM2 may mediate a protective effect in neurons under excitotoxic conditions^120^. Whether phenethylaminylation or other psychedelic protein modifications could enhance such an effect would constitute a novel area of investigation.

### Extracellular Matrix (ECM) Proteins

Many of the initial studies on monoaminylation examined the modification of fibronectin, a major glycoprotein important to the maintenance of both structure and signaling in the ECM^34, 121^. In the CNS, *in vitro* evidence suggests the possibility that fibronectin serotonylation by glial cells increases the deposition of ECM proteins, constituting a potentially pro-fibrotic mechanism^34^. In the periphery, fibronectin serotonylation by a competing transglutaminase, Factor XIII-A, decreases collagen fibrillogenesis, constituting a potentially anti-fibrotic mechanism^39^. Fibronectin serotonylation also promotes the proliferation and migration of smooth muscle cells, suggesting that such fibronectin modifications are important in the cellular response to ECM signals^38, 122–124^. Brain ECM reorganization is thought to be an important mechanism by which psychedelics, particularly psilocybin and 3,4-methylenedioxymethamphetamine (MDMA), “reopen” critical periods in mature animals, reenabling social reward learning^125, 126^. Enduring ECM protein modifications by such tryptamines and phenethylamines could, hypothetically, promote long-term ECM remodeling.

### Ribosomal Proteins

The 40S ribosomal protein S19 has previously been reported to be serotonylated^72^. However, the effect of ribosome protein monoaminylation is unknown. There is evidence that TGM2-mediated polyamination of eukaryotic translation initiation factor 4E-binding proteins preferentially enhances the translation of mRNAs containing G/C-rich 5’-untranslated regions^127^. In a similar manner, perhaps the modification of proteins involved in ribosome-mRNA interactions by psychedelics could shift the translatome of cells.

### Histones

The potential for psychedelics to act as histone marks is intriguing given the compelling works of Ian Maze’s group robustly identifying H3 serotonylation, dopaminylation and histaminylation^128^. Likely of major interest to psychedelic researchers are the findings that these histone marks are involved in the function of critical brain regions such as the dorsal raphe and ventral tegmental area, where they contribute to both the pathophysiology of neuropsychiatric disorders and the therapeutic mechanisms of their current treatments, as previously discussed^66, 67^.

Research on protein modifications by psychedelics can move forward in several different directions, each with the potential to improve our understanding of how psychedelics modulate cellular processes of the brain. In our pilot study, we demonstrate that a diverse set of proteins are potential targets of TGM2-mediated phenethylaminylation by 4PM, an alkyne-functionalized derivative of mescaline, in primary human astrocytes. In doing so, we highlight 4PM as a useful tool for preliminary investigations of monoaminylation by psychedelics utilizing click-chemistry techniques. At a small scale, our simplified synthetic route to produce 4PM may be sufficient and convenient for basic scientists studying the cellular and molecular mechanisms of protein modifications by psychedelics. Even if direct modification of proteins by psychedelic compounds does not occur appreciably *in vivo*, it may be worth studying how psychedelic signaling pathways modulate the TGM2-mediated modification of proteins by endogenous monoamines. With continued examination of these mechanisms, we may better elucidate the significance, if any, of the protein psychedelome.

## Methods

### Cell Culture

Primary human optic nerve head (ONH) astrocytes were used for *in vitro* experiments. ONH astrocytes were utilized due to the accessibility of tissues from the anterior visual system and the relative ease with which mature astrocytes from the ONH can be cultured – continuous culture of post-mitotic glial cells isolated from post-mortem human brains is currently a challenge^129^. ONH astrocytes broadly resemble other fibrous white matter astrocyte populations found throughout the CNS and are a *TGM2*^+^ cell type^130–132^. Isolation and culture of ONH astrocytes was performed using previously described methods^133, 134^. In brief, human donor eyes were obtained from the UNTHSC Center for Anatomical Sciences. Eyes were collected within 24 hours post-mortem and managed with adherence to the Declaration of Helsinki. Optic nerves were separated from the eyes and the unmyelinated ONH region was carefully dissected^135^. ONH explant tissues were grown on 12-well plates with Ham’s F10 medium supplemented with 10% FBS, 0.3 mg/mL L-glutamine and 1% penicillin/streptomycin in a humidified incubator (37 °C, 5% CO_2_) until cells migrated out and reached confluence. Astrocytes were selectively grown from the mixed ONH cell population using 72 hours of serum-deprivation. Once isolated, ONH astrocytes were maintained in astrocyte basal medium supplemented with 5% FBS and 1% penicillin/streptomycin (ScienCell, Carlsbad, CA). Three biologically distinct cell strains from different human donors were used in every following experiment (n=3). Cell characterization was performed as previously described by immunocytological detection of glial fibrillary acidic protein (GFAP) and neural cell adhesion molecule 1 (NCAM/CD56) (Fig. 9)^133, 136^.

**Figure 9.**
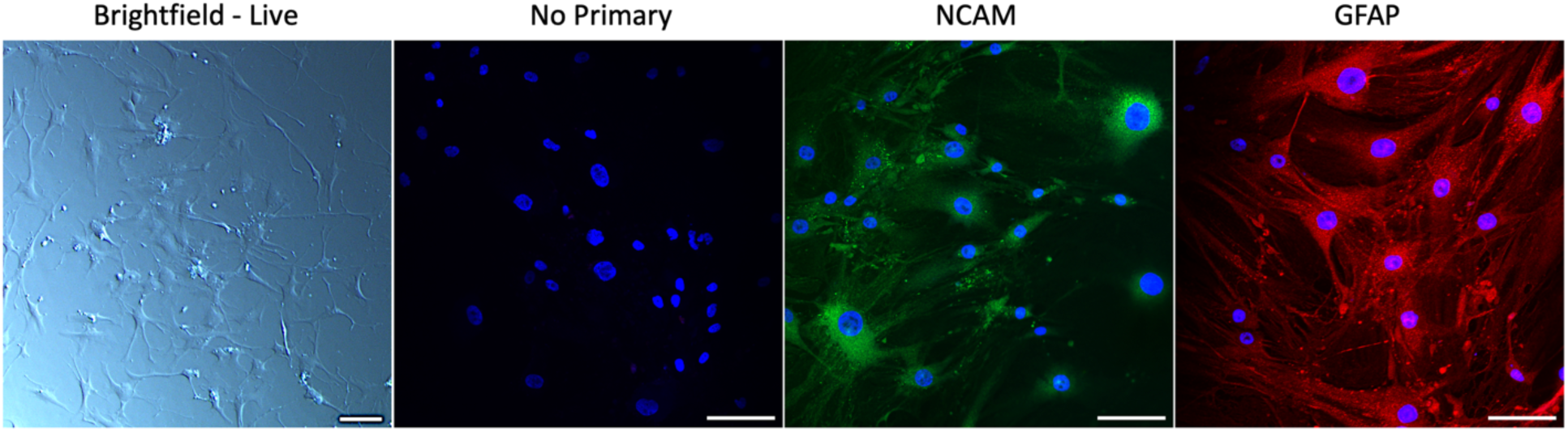
Characterization of primary human ONH astrocytes. From left to right: live brightfield image of cells displaying typical astrocytic morphology (image uniformly brightened and sharpened to improve visibility of processes, scale bar = 100 μm); immunofluorescent image of ONH astrocytes stained with DAPI and either no primary antibody, NCAM (green), or GFAP (red) (Leica DMi8 Confocal, total magnification = 200x, scale bar = 100 μm).

### Click-Chemistry Detection of 3,5-Dimethoxy-4-(2-Propynyloxy)-Phenethylamine in Fixed ONH Astrocytes

Primary human ONH astrocytes were grown to confluence on 24-well plates and then serum-deprived for 24 hours. Cells were then treated with either 0, 1, 50, 100, 500 or 1000 μM of 3,5-dimethoxy-4-(2-propynyloxy)-phenethylamine hydrochloride. After 24 hours, cells were gently washed, fixed in 4% paraformaldehyde, and permeabilized with 0.2% TritonX-100. A biotinylated azide (PEG4 carboxamide-6-azidohexanyl biotin, 5 μM) was “clicked” onto the alkyne moiety of intracellular 3,5-dimethoxy-4-(2-propynyloxy)-phenethylamine molecules through the CuAAC reaction using a Click-iT cell reaction buffer kit following manufacturer instructions (C10269, Thermofisher). Biotin tags were then labeled with AlexaFluor 647 fluorophore-conjugated streptavidin (1:500). Nuclei were counterstained with Hoechst and cells were imaged with a Keyence BZ-X710. Fluorescence intensity was calculated as mean gray value (after auto-thresholding) and expressed as fold-changes relative to untreated controls. Data normality was inferred by a Shapiro-Wilk test. Significance was calculated by a repeated measures ANOVA (GraphPad Prism, multiple comparisons analyzed using Fisher’s LSD test, biological n=3, significance defined as p<0.05).

### Calcium Assays

To assess the ability of 3,5-Dimethoxy-4-(2-Propynyloxy)-Phenethylamine to induce intracellular Ca^2+^ elevation, a qualitative fluorescence microscopy assay using Fluo-4 was utilized. Fluo-4 is a 1,2-Bis(2-aminophenoxy)ethane-N,N,N’,N’-tetraacetic acid (BAPTA) based calcium indicator which exhibits increased green fluorescence intensity upon calcium binding. Primary human ONH astrocytes (n=3) were grown to confluence on 24-well plates and then serum-deprived for 24 hours. All cells were then incubated with 2.5 μM of cell permeant Fluo-4 acetoxymethyl ester (AM) (Thermofisher) prepared in a 1:1 solution of DMSO and Pluronic F-127 for 30 minutes at 37 °C followed by 15 minutes at room temperature to allow for cellular uptake and de-esterification of the calcium indicator. 10 μM of BAPTA AM (Thermofisher) was similarly prepared and delivered to negative control cells to chelate intracellular Ca^2+^ simultaneously. After incubation, cells were washed with PBS and then kept in phenol-free cell culture media for imaging using a Keyence BZ-X710. Images were taken first at baseline, and then at 60 seconds following treatment with 100 μM 3,5-Dimethoxy-4-(2-Propynyloxy)-Phenethylamine, 100 μM serotonin (5-HT) or 2.5 μM Calcimycin (calcium ionophore A23187; positive control). The media of untreated cells was mixed by gentle pipetting to simulate compound addition.

To provide a quantitative validation of the Fluo-4 calcium assay, a Fura-2 calcium assay was performed. Fura-2 is a fluorescent calcium indicator that always exhibits peak emission at around 510 nm but can be excited at either 340 nm or 380 nm. The excitation maximum of Fura-2 shifts in the presence of calcium, enabling ratiometric measurements of fluorescence intensity^137, 138^. Similar to above, confluent cells grown on 8-well Nunc Lab-Tek II chambered coverglass were serum deprived and then incubated with Fura-2 AM (Thermofisher) for 30 minutes at 37 °C before being washed and imaged. Fura-2 imaging was performed using an Olympus IX81 inverted fluorescence microscope. Measurements were recorded and analyzed using MetaFluor (Molecular Devices, San Jose, CA), which automatically records 510 nm fluorescence emission intensity after 340 nm excitation and 380 nm excitation to generate F_340_/F_380_ ratios. Measurements were recorded for individual cells, with several cells independently analyzed for each biological replicate. Recordings of Fura-2 F_340_/F_380_ ratio values were taken every second, with 5 baseline measurements followed by 240 measurements after the addition of 100 μM 4PM. Data points from every fifth second were plotted to visualize the effect of 4PM treatment on intracellular Ca^2+^ levels over time.

### Click-Chemistry Detection of 3,5-Dimethoxy-4-(2-Propynyloxy)-Phenethylamine in ONH Astrocyte Protein Lysates

Primary human ONH astrocytes were grown to confluence on 6-well plates. Cells were then pre-treated with either a vehicle solution (DMEM with 10% serum and 2% DMSO) or 100 μM Z-DON for 2 hours. Z-DON is a peptide (Z-QVPL) based irreversible inhibitor of transglutaminase activity with selectivity for TGM2^139–141^. After the pre-treatment period, cells were stimulated by the addition of 1.5 μM Calcimycin (to provide the high Ca^2+^ cellular environment needed for TGM2 activity) and incubated with 100 μM 3,5-Dimethoxy-4-(2-Propynyloxy)-Phenethylamine for 6 hours (a shorter timepoint was used here to minimize cytotoxicity from Calcimycin). Cells were then washed 3 times with PBS and lysed in ice cold MPER lysis buffer with Halt protease inhibitor. Protein lysates were spun down at 10000 RCF for 10 minutes to remove cell debris. The lysates were then incubated with Pierce streptavidin magnetic beads (Thermofisher) for 1 hour at room temperature on an end-over-end rotating platform to clear the lysates of any endogenously biotinylated proteins^142^. Protein was re-extracted from the pre-cleared lysates using a methanol:chloroform:water precipitation to wash out any free drug remaining in the lysates. This step was taken to reduce the formation of nonspecific thiotriazoles during the proceeding click chemistry reaction – thiotriazoles can form from the reaction of free alkyne probes with thiols on proteins and the azide tag, and are difficult to distinguish from real *in cellulo* labeling^143^. Protein samples were reconstituted in 50 mM TBS with 0.5% SDS (pH=7.4) and treated with 50 mM iodoacetamide for 30 minutes to alkylate cysteine residues, reducing the availability of reactive thiols for nonspecific binding by the biotin-azide tag^144^. Biotin tags (biotin-azide 5 μM) were “clicked” onto the alkyne moiety of phenethylamine-modified cellular proteins (50 μg) in our samples through a CuAAC reaction using a Click-iT cell reaction buffer kit (C10276, Thermofisher). As a slight modification to the kit’s manufacturer instructions, 1 mM L-cysteine hydrochloride monohydrate was added to the reaction mixtures to reduce the ability of nonspecific thiotriazole intermediates to react with proteins^143^. The reaction mixtures were cleaned and proteins extracted through a standard methanol:chloroform:water precipitation procedure. The resultant protein pellets were air-dried under sterile airflow. Proteins were reconstituted in MPER protein buffer with Halt protease inhibitor and incubated with streptavidin magnetic beads for 1 hour at room temperature on an end-over-end rotating platform. Beads were then washed and the enriched biotinylated protein samples were eluted by heating at 98 °C for 15 minutes in SDS denaturing sample buffer. The samples were separated by SDS-PAGE on 4-12% precast bis-tris polyacrylamide gels for western blotting and mass spectrometry analysis.

For immunoblotting, proteins from the gels were transferred onto a PVDF membrane, blocked for 1 hour in SuperBlock blocking buffer (Thermofisher) and then incubated overnight at 4 °C with an anti-biotin antibody (Thermofisher 31852, 1:1000). The blot was washed and incubated in an appropriate HRP-conjugated secondary antibody (Thermofisher, 1:5000) for 1 hour at room temperature and imaged via chemiluminescence with a ChemiDoc imaging system (BioRad, Hercules, CA). All distinct bands were quantified via integrated density analysis, with the exception of a prominent 15 kDa band representing leeched streptavidin monomers. The sums of integrated densities were normalized against GAPDH from clicked, but unenriched, cellular lysates. Data normality was inferred by a Shapiro-Wilk test. Significance was calculated by a repeated measures ANOVA (GraphPad Prism, multiple comparisons analyzed using Tukey’s HSD test, biological n=3, significance defined as p<0.05).

For gel analysis, polyacrylamide gels were fixed using a mixture of acetic acid, methanol and water and then stained with GelCode Blue stain reagent (Thermofisher). Gel bands from 3,5-Dimethoxy-4-(2-Propynyloxy)-Phenethylamine-treated samples were excised and sent to the University of Texas Southwestern Proteomics Core for protein identification via 90-minute reverse-phase LC-MS/MS (QExactive HF) followed by Proteome Discoverer analysis. The top 150 abundant proteins were used to generate a string protein-protein physical interaction network, which was organized by major function or cellular localization.

## Acknowledgements

We are grateful to Jennifer Pham, Amanda Tate, Pinkal Patel, Stacy Curry, Raghu Krishnamoorthy, Denise Inman and Nathalie Sumien for their assistance and support. We would like to thank Andrew Lemoff and the University of Texas Southwestern Proteomics Core, as well as Donna Coyle, Carlos Castello and the University of North Texas Health Science Center Pharmaceutical Analysis Core Lab for their technical assistance. Lastly, we would like to express our sincere appreciation for the human tissue donors and their invaluable gift, which enabled this study.

Figures 1 & 2 were created with BioRender.com.

This study was supported by a NIH National Institute on Aging training grant (T32 AG020494).

